# *Compartmap* enables inference of higher-order chromatin structure in individual cells from scRNA-seq and scATAC-seq

**DOI:** 10.1101/2021.05.17.444465

**Authors:** Benjamin K. Johnson, Jean-Philippe Fortin, Kasper D. Hansen, Hui Shen, Timothy Triche

## Abstract

Single-cell profiling of chromatin structure remains a challenge due to cost, throughput, and resolution. We introduce *compartmap* to reconstruct higher-order chromatin domains in individual cells from transcriptomic (RNAseq) and epigenomic (ATACseq) assays. In cell lines and primary human samples, *compartmap* infers higher-order chromatin structure comparable to specialized chromatin capture methods, and identifies clinically relevant structural alterations in single cells. This provides a common lens to integrate transcriptional and epigenomic results, linking higher-order chromatin architecture to gene regulation and to clinically relevant phenotypes in individual cells.

## Introduction

Mammalian genomes are often presented as linear sequences, one per chromosome. A linear depiction emphasizes the impact of base substitutions, deletions, and duplications in shaping life as we know it. This can obscure another remarkable feature of DNA: its robustness to three-dimensional manipulation. Meters of DNA, and all of its associated machinery, are compressed to fit into a cell nucleus only micrometers across^1^. Cells must access the right sets of genetic instructions at the right time in the right cells for development to succeed^2^. The emergence of complex multicellular organisms has, in part, mirrored the emergence of progressively refined mechanisms that pack, unpack, and transcribe DNA^3–8^. The three-dimensional structure of the genome in living cells is thus no less central to multicellular life than the sequence of bases that comprise DNA.

Unpacking DNA from chromatin fibers exposes “beads on a string”: nucleosomal beads slide along a DNA string to permit assembly of protein complexes ^9–11^. Some of these, such as cohesin and condensin, seem to function primarily in structural roles. Others, such as the pre-initiation complex, primarily enable RNA transcription. Yet others, such as the massive Mediator and Integrator complexes, span both roles ^12–20^. Recent work in developmental and cancer biology has revealed that alterations in structural complexes can impact transcription in ways unrelated to genome integrity, causing developmental abnormalities and premalignant conditions ^21–31^. In otherwise healthy organisms, viral transcriptional read-through can remodel the physical arrangement of surrounding DNA. The rapid advance of single-cell genomic assays has permitted inspection of these and other fundamental phenomena in individual cells from primary tissues ^29,30,32-45^.

Many recent advances in understanding 3D chromatin architecture have come from conformation capture methods, whether ligation-dependent (e.g., Hi-C) or independent (e.g., SPRITE), and via *in situ* imaging ^46,47^. Traditional conformation capture methods require thousands of cells and deep sequencing to distinguish chromatin contacts from background signal ^16^. These requirements created technical and logistical challenges, limiting genome-scale use to experiments where large numbers of putatively similar cells could be obtained. Super-resolution imaging approaches are inherently suited for single cell investigations, but are limited in throughput, and require specialized instruments ^34,37,40^. Newer conformation capture methods reduce input and sequencing requirements, with the goal of dissecting chromatin folding in single cells ^46,47^. These approaches trade information density for throughput, a pervasive phenomenon in single-cell assays not limited to chromatin capture methods or microscopy^48^. A complementary approach is to impute missing data, and indeed, several computational approaches have been developed to denoise chromatin contact information ^49–55^.

Gene-rich, actively transcribed, megabase-scale regions in the euchromatic “open” or “A” compartment of chromosomes typically localize to the chromosomal surface. The corresponding “closed” or “B” compartment is generally enriched for heterochromatin. At finer scales, structures referred to as topologically associating domains (TADs) have varied roles in directing specific gene regulatory programs, with the maintenance of TAD boundaries remaining an open research question ^3^. In mammals, these boundaries are enriched for CTCF binding sites that appear to recruit machinery that maintains defined borders. Disrupted boundaries can promote developmental defects or pro-malignant states due to loss of temporal and cell-type-specific gene regulatory programs ^3^.

Chromatin architecture is intertwined with transcriptional and epigenetic regulatory mechanisms. Thus, another approach to address limitations of specialized approaches employs orthogonal single-cell assays, such as single-cell whole-genome bisulfite sequencing (scWGBS), single-cell ATAC-seq (scATAC-seq), and scRNA-seq. In recent years, methods have emerged to infer global and focal chromatin architecture in groups of samples as small as a few cells ^56–60^. Inference of chromatin structure from common single-cell assays can yield functional readouts absent from more specialized assays.

Recent work has shown that A/B compartment and TAD(-like) structure boundaries defined by bulk conformation capture methods are dynamic in single cells, an emergent property of modeling of many cells ^37,61^. This is consistent with the heterogeneity seen in single-cell gene expression experiments, given that structural boundaries of chromatin domains can influence nearby gene expression. Further, a recently published method, RD-SPRITE, demonstrated that RNA can modify higher-order chromatin interactions involved in essential nuclear functions ^27^. Single-cell methods can provide insight into the structural basis of chromatin changes by integrating genome-wide DNA and RNA measurements, providing context and a more complete picture of cellular state & fate.

Here, we capitalize upon these features, using an empirical Bayesian approach to extend previous work to single-cell inference, and to infer transcriptional consequences in development, disease, and disorder of individual cells ^56^. Our methods, available in the Bioconductor package *compartmap*, provide a unifying approach to probe chromatin structure, regulation, and dynamics via common bulk and single-cell assays, such as single-cell RNA sequencing (scRNA-seq). We show that the close relationships between transcription, accessibility, and genome structure make inference of higher-order chromatin structure feasible from transcriptional measurements alone. We generalize previous work showing that correlated changes in accessibility and DNA methylation can recover three-dimensional chromatin structures similar to Hi-C. Further, we demonstrate that features of chromatin domains captured by transcription are both comparable and complementary to those observed by chromatin capture techniques.

## Results

Previous work has demonstrated that position-dependent and -independent^62^ effects on transcription could be separated in bulk RNAseq data, while both DNA methylation and accessibility (ATACseq) assays could reconstruct chromatin compartments as observed via Hi-C^56^. We reasoned that shrinkage and resampling could extend uncertainty-aware inference of chromatin structure from any of these assays to the level of individual cells.

*Compartmap* builds on the preceding, and follows four steps:

1. preprocessing and summarizing the single-cell assay data,
2. empirical Bayes shrinkage of summarized features towards a global or local target,
3. computing the shrinkage correlation estimate of summarized features from (2), and
4. computing higher-order domains via singular value decomposition (SVD) (**Figure 1a**).

**Figure 1.**
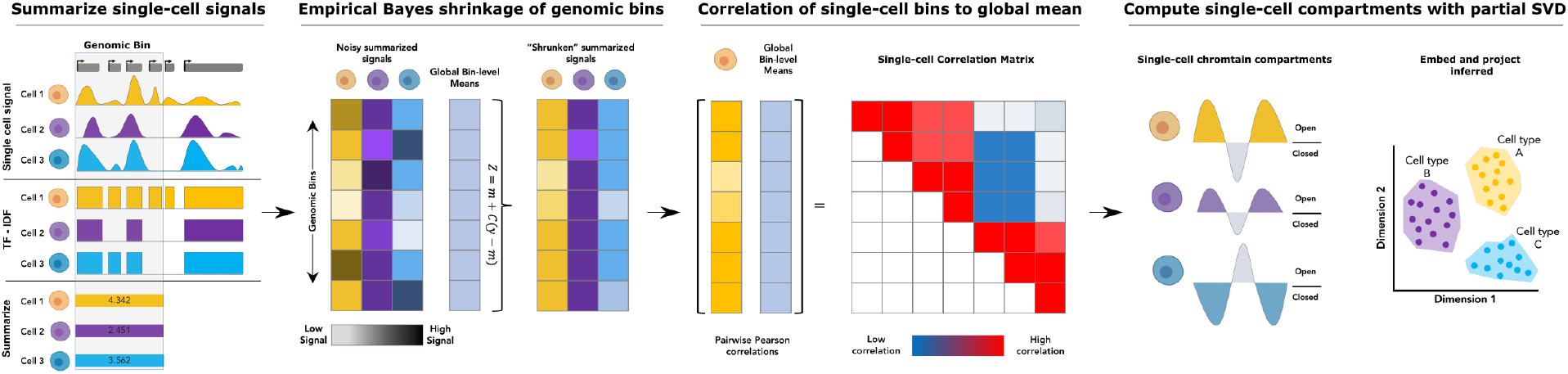
Compartmap reconstructs higher-order chromatin structures in single cells. The *compartmap* workflow performs term-frequency inverse document frequency (TF-IDF) transforms and summarizes single-cell signals into genomic bins (e.g., 100 kb). Summarized signals are shrunk towards a global or targeted mean using an empirical Bayes approach. Pairwise correlations between the single-cell shrunken bins and the global or targeted mean are computed, followed by partial singular value decomposition (SVD). Group- or population-level compartments can also be computed from correlations of shrunken single-cell signals across all cells present in the matrix as opposed to the global mean. The first right singular vector is retained, which is associated with open and closed chromatin compartments. Positive eigenvalues correspond to euchromatic or A compartments, while negative eigenvalues correspond to heterochromatic or B compartments.

In (1), we binarize single-cell signals (e.g., gene expression) and apply term-frequency inverse-document frequency (TF-IDF) transformation. The transformed features are then summarized into genomic bins of arbitrary size (default 100 kb) (see Methods). In (2), binned, summarized features are shrunken towards a targeted mean using a James-Stein estimator (**Equation 3** in Materials and Methods). Note that a reference sample or population can be targeted, as when comparing changes in response to a treatment or within a cell type, allowing for targeted shrinkage approaches as desired. In (3), we construct a pairwise correlation matrix of shrunken genomic bins for each cell. In (4), we perform SVD of the single-cell correlation matrix **M** into **UΣV^T^**, retaining the first right singular vector (**V**_1_, the first column of **V**). The latter recalls Hi-C approaches to infer A/B compartments, where the first principal component represents open & closed domains. In *compartmap*, we also smooth **V**_1_ with a sliding window approach (Materials and Methods), and implement a novel resampling-based scheme to capture uncertainty regarding the sign and changepoints of entries in **V**_1_, as described in more detail below.

Due to the sparsity of single-cell data, we employ a bootstrap procedure to quantify uncertainty about inferred compartments ^63^. *B* bootstrap genome-wide global means are drawn from the data. Following initial computation of domains in a cell, the procedure is repeated for each bootstrap global mean. Results are tallied as “open” or “closed” per the sign of the first right singular vector **V**_1_, computing Agresti-Coull confidence intervals on each bin ^64^. For a real symmetric matrix, the SVD is unique up to unit transformation on each column of **U** and **V**. “Sign flips” are thus an inevitable consequence, and are also observed in Hi-C. By fully propagating uncertainty, we enable users to rapidly identify and correct “sign flips.” As a side effect, this also allows pooling of information across adjacent bins and similar cells, weighted by uncertainty, enabling multi-scale inference on boundaries of compartments and similar structures across cells.

### Compartmap identifies single-cell domains without proximity ligation

As a proof of concept, we applied *compartmap* to the human erythroleukemic cell line K562. The Tier 1 status of K562 within the ENCODE and 4DN consortia, and its use as a model system elsewhere, has generated a wealth of existing data, including high-resolution Hi-C and scHi-C data ^16,32,65,66^. Given the level of aneuploidy observed in K562, we first looked at chromosome 14, highlighted in prior work as diploid in this cell line ^16,56,66^. Chromatin compartments were computed at 100-kb resolution for Hi-C, single-cell combinatorial indexing Hi-C (sci-Hi-C), and single-cell ribo-reduced total RNA-seq (scRNA) data. Qualitatively, sci-Hi-C- and *compartmap*-inferred domains are consistent in aggregate and in single cells, relative to bulk Hi-C (gray shaded boxes, **Figure 2a**). However, there exist regions where *compartmap*-inferred chromatin domains deviate from those of sci-Hi-C and bulk Hi-C (red shaded boxes, **Figure 2a**). To explore whether euchromatic regions (as inferred by *compartmap*), nested within heterochromatic regions (as called from single-cell and bulk Hi-C data) were artifactual, we evaluated orthogonal DNase hypersensitivity-seq (DHS) and Lamin A/B/C TSA-seq data across these regions. Notably, where *compartmap* inferred “open” chromatin states from scRNA among “closed” regions from Hi-C data, the presence of DHS signal and absence of Lamin A/C TSA-seq signal ^67^ corroborate the *compartmap* calls. Where fine-scale chromatin domains are inferred by *compartmap* and distinct from sci-Hi-C or bulk Hi-C, they are also supported by orthogonal chromatin structure measurements. *In situ* genome sequencing^68^ has recently highlighted the role of such single-cell domains.

**Figure 2.**
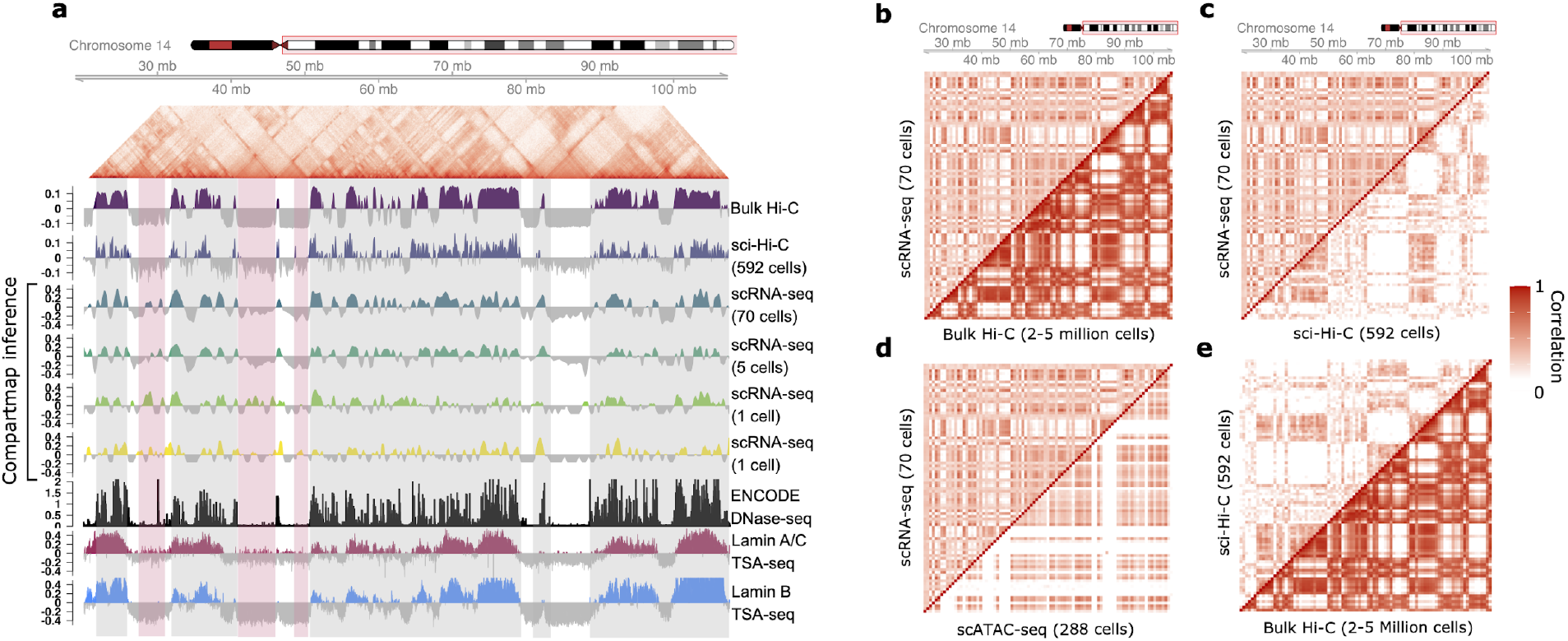
*Compartmap* reconstructs chromatin domains across multiple genomic scales in single cells. (a) *Compartmap* can reconstruct chromatin domains from riboreduced single-cell total scRNA-seq in K562 cells, similar to bulk Hi-C and aggregate sci-Hi-C at 100-kb resolution (chromosome 14). Gray boxes highlight similar chromatin domain structures, while red boxes denote deviations from chromatin-capture-based methods relative to the scRNA-seq-inferred domains at the given resolution. Inferred scRNA-seq chromatin domain deviations are supported by apparent substructure in the bulk Hi-C contact matrix (top), accessibility from ENCODE DNase-seq (black track), and Lamin A/B/C TSA-seq (red and blue tracks) as orthogonal data. Lamin A/B/C tracks are inverted where red/blue colors represent non-Lamin associated domains. Positive eigenvalues correspond to “open” chromatin domains and negative eigenvalues correspond to “closed” chromatin domains. (b) Comparison of group level correlation matrices of chromosome 14 at 1-mb resolution show similar resolution and structure between riboreduced single-cell total scRNA-seq (*N* = 70 cells) and bulk Hi-C (*N* = 2-5 million cells), (c) total scRNA-seq (*N* = 70 cells) and sci-Hi-C (*N* = 592 cells), (d) total scRNA-seq (*N* = 70 cells) and scATAC-seq (288 cells), and (e) sci-Hi-C (*N* = 592 cells) and bulk Hi-C (*N* = 2-5 million cells). scRNA-seq and scATAC-seq correlation matrices were generated using *compartmap*. Bulk Hi-C and sci-Hi-C correlation matrices were computed from normalized contact matrices. Coloration represents low (white) to high (red) within-platform correlations.

Chromatin contact matrices provide a rich view of the multi-scale nature of chromatin organization. Within the context of chromatin-capture-based methods, prior work has shown that by binning and computing correlation matrices at larger genomic scales (e.g., 1 Mb, megabase resolution) from the normalized contact matrix, a higher resolution view of interacting domains can be observed, via a plaid-like patterning ^69^. To assess whether the correlation matrices produced via *compartmap* from the K562 scRNA-seq and scATAC-seq correspond to those observed by bulk and sci-Hi-C, we overlaid aggregate single-cell profiles with those from the chromatin-capture-based methods (**Figure 2b-e**). Strikingly, at a resolution of 1Mb, a similar plaid-like patterning was observed between aggregate scRNA-seq (*N* = 70 cells) and bulk Hi-C (*N* = 2-5 million cells), supporting prior observations at smaller genomic scales (100 kb) that chromatin domains can be inferred from scRNA-seq, similar to bulk and single-cell chromatin-capture-based methods (**Figure 2a-b**). Additionally, this supports the well-established links between gene expression and chromatin architecture, and also demonstrates the high density of information contained within total single-cell transcriptional profiling approaches. Further, *compartmap* is able to reconstruct these plaid-like domains in other cell types and single-cell sequencing approaches, suggesting that the observed results in K562 are not unique to this cell type or single-cell sequencing technologies (**Figure S5**). Notably, ribo-reduced total scRNA-seq approaches (e.g., STORM-seq and SMART-seq total) produce both group-level correlation and inferred higher-order chromatin domains that are more similar to bulk Hi-C measurements than to poly-A selected protocols (**Figure 2b and S5**).

The scRNA-seq correlation matrix patterning in K562 was comparable with that of aggregate sci-Hi-C, though the overall sparsity of sci-Hi-C is higher at this genomic scale relative to either the scRNA-seq or bulk Hi-C (**Figure 2c,e**). Intriguingly, when the scRNA-seq and scATAC-seq correlation matrices were overlaid, an overall similar substructure in the patterning was observed (**Figure 2d**). This result corroborates prior work with which *compartmap* is built upon, showing that higher-order chromatin can be inferred from scATAC-seq and other, non-chromatin-capture-based methods such as methylation arrays ^56^. However, to our knowledge, this is the first demonstration that scRNA-seq and scATAC-seq can directly provide similar chromatin domain architecture information, giving a physical lens via higher-order chromatin to integrate orthogonal single-cell transcription and accessibility datasets in individual cells.

### Compartmap boundaries are enriched for CTCF sites

Next, to quantitatively determine whether the inferred compartment boundaries were enriched for CTCF sites similar to chromatin conformation capture methods, we computed the distance to the nearest CTCF site in each data type. As expected, both Hi-C and sci-Hi-C compartment boundaries are close to annotated CTCF sites (less than 25 kb), both within chromosome 14 and genome-wide (**Figure S2a**)^16,32^. Similarly, the compartment boundaries inferred by *compartmap* in both aggregate and single cells have similar distances to CTCF sites, within and across the genome (**Figure S2a**). Taken together, these results provide evidence that *compartmap* is able to infer chromatin domains with biologically-relevant features, in aggregate and in single cells using scRNA-seq, recovering comparable information to traditional proximity-ligation and chromatin conformation capture approaches, without need for specialized assays.

We further investigated bin-level (100-kb) correlations of aggregate sci-Hi-C and *compartmap* calls from scRNA-seq to bulk Hi-C. In aggregate, *compartmap* and sci-Hi-C exhibit similar correlations across chromosomes to bulk Hi-C data with overall mean Pearson correlations close to *r* = 0.75 (**Figure S2b**). As expected, when single-cell correlations were computed against bulk Hi-C, the genome-wide correlations dropped to approximately *r* = 0.55 (**Figure S2b**). These results are consistent with prior observations that single-cell chromatin domains are more dynamic than those observed in population averaged Hi-C, but still maintain boundaries that are enriched for nearby CTCF sites (**Figure S2a**)^37^. Indeed, a random subset of single cells taken from the larger K562 scRNA-seq dataset (*N* = 70; **Figure 2a**), when aggregated together, reconstruct similar features as those observed in bulk Hi-C at larger genomic scales (1-mb resolution) with more heterogeneous compartment structures observed in individual cells (**Figure S3a**), and maintain domain boundaries with coherent distances to nearby CTCF sites (**Figure S3b**).

To identify whether the heterogeneous chromatin structures observed in individual cells were due to variable compartment estimates, we calculated bootstrap confidence estimates for each cell across the genome (**Figure S4**). The bulk of single-cell bootstrap confidence estimates exceeded 80% sign coherence, suggesting that the observed heterogeneity in single-cell chromatin domains likely reflects underlying biology (**Figure S4**). Cell cycle heterogeneity has previously been observed in single-cell higher-order chromatin structure^70^, so we applied several approaches to infer single-cell cycle states from scRNA-seq. We found all cells classified as S or G2/M phase, where previous work has indicated that chromatin structures are largely stable^70,71^. We take this to indicate that, consistent with recent *in situ* genome sequencing results ^68^ and previous microscopic examinations, the cell-private domains inferred by *compartmap* represent biological (rather than technical) variability. Thus *compartmap* extracts orthogonal information from popular single-cell assays without increasing their associated cost or difficulty. When the assay is scRNA-seq, *compartmap* complements the results of proximity ligation assays by providing direct transcriptional readouts.

### Compartmap detects complex structural variants lacking fusion transcripts

We sought to determine if *compartmap* could provide orthogonal benefit to WGS and traditional analyses of bulk or single-cell RNA sequencing in a translational context. Short-read whole-genome sequencing (WGS) has clear clinical benefits^72^, but validation of novel structural variants (SVs), clonal history, and their consequences is challenging. Chromatin capture methods can identify complex SVs via off-diagonal correlations, but input cell requirements can be prohibitive ^73,74^, and transcriptional consequences unclear. RNA sequencing sensitively detects rare fusion gene transcripts, but requires filtering for *trans*-splicing events and validation of results. “Enhancer hijacking”^73,75^ SVs lack fusion transcripts, further complicating matters. This mechanism is seen in the recurrent inv(3)(q21;q26) of aggressive myelodysplastic syndromes (MDS) and acute myeloid leukemia (AML). Gröschel and colleagues found inv(3)(q21;q26) translocates a strong *GATA2* enhancer (3q21) into the promoter of the *EVI1 (MECOM)* gene (3q26). This deactivates a key transcription factor, activates a powerful oncogene, and yields dismal outcomes: only 3% of patients survive 5 years^76^. Inv(3) patients resist standard induction chemotherapy, thus identification of 3q involvement has high diagnostic yield.

To assess whether *compartmap* can identify disrupted chromatin domains in primary clinical samples, we reanalyzed scRNA-seq data from the unsorted bone marrow aspirates of an inv(3)(q21;q26) AML patient^77^. The correlation matrices for chromosome 3 at 1 Mb resolution were broadly similar in normal and malignant cells (**Figure 3a**). However, taking the difference of the two cell-type correlation matrices readily revealed the chromatin structures disrupted by the inversion, with interaction breakpoints proximal to *GATA2* and *MECOM* (**Figure 3b-c**). Of note, in cells with inv(3)(q21;q26), ectopic expression of *EVI1* also leads to expansion of the closed “B” compartment, as shown in **Figure 3c**, consistent with EVI1’s orchestration of H3K9 methyltransferase activity, heterochromatin formation, self-renewal, and treatment resistance. ^78–80^ *Compartmap* inference of chromatin disruption in primary clinical samples can thus help prioritize and filter SVs, which are difficult to interpret or validate from RNA-seq alone. Even in patients with whole-genome sequencing or centralized metaphase karyotyping, the presence of multiple and/or cryptic SVs presents a formidable diagnostic challenge. This proof of principle shows how *compartmap* can help to address these challenges.

**Figure 3.**
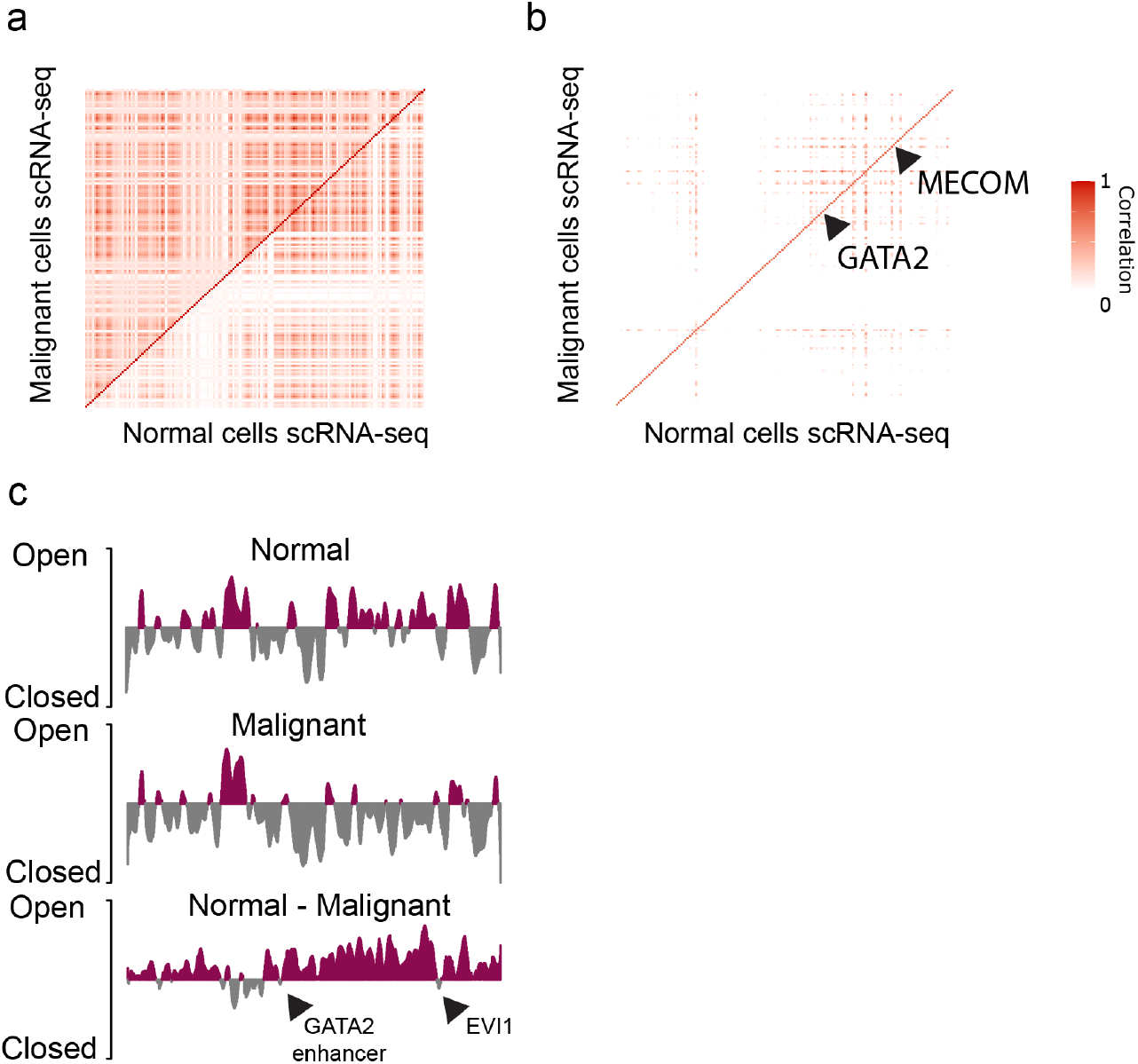
*Compartmap* can identify global chromatin disruption due to enhancer hijacking from scRNA-seq in primary inv(3)(q21;q26) AML. (a) De-noised correlation matrices of higher-order chromatin interactions on chromosome 3 at 1-mb resolution in malignant and normal scRNA-seq of a single patient with inv(3) AML. (b) Difference plot of normal and malignant correlation matrices demonstrate the altered chromatin interactions due to inv(3) on the q arm of chromosome 3, corresponding to relocating the *GATA2* enhancer to the *MECOM* locus. Colors represent low (white) to high (red) within-platform correlations. (c) *Compartmap*-inferred higher-order chromatin domains at 1-mb resolution of chromosome 3 of normal, malignant, and the difference supports the clonal SV inv(3), leading to altered chromatin interactions with two key differential domains containing the *GATA2* enhancer and *EVI1* locus.

## Discussion

Higher-order chromatin structure plays critical roles during development and disease. Disruption of the chromatin regulatory and maintenance machinery can lead to negative phenotypic effects at the cellular, tissue, and whole-organism levels. Thus, many methods have been developed to either directly assay or infer chromatin states from a variety of cell and tissue sources. With the introduction of single-cell approaches to probe chromatin architecture, a far more dynamic and heterogeneous view emerges, corroborating and challenging existing dogma in the field that was largely derived from population-averaged or bulk chromatin conformation capture experiments.

Here, we demonstrate a complementary approach for inferring higher-order chromatin structure in single cells from single-cell RNA-seq and single-cell epigenetic assays. As a proof-of-concept, the application of *compartmap* to K562 total single-cell RNA-seq was able to reconstruct population averaged/bulk high-depth Hi-C and sci-Hi-C data in the same cell line. Additionally, the aggregate total scRNA-seq (70 cells) and aggregate sci-Hi-C (approximately 500 cells) produced similar chromatin domain architecture, qualitatively and quantitatively across the genome, when compared with bulk Hi-C data. Notably, *compartmap*-inferred chromatin domain deviations from both proximity ligation techniques (Hi-C and sci-Hi-C) in aggregate and single cells is supported by both domain boundary distances to CTCF sites and other measures of chromatin accessibility, i.e., DNase-seq and TSA-seq (**Figure S2, 2a**). Thus, it is tempting to speculate that the observed single-cell chromatin dynamics inferred by *compartmap* in K562 may be real and not an artifact of the inference procedure. However, we note that the observed continuous signal across the genome in the total scRNA-seq used as a proof-of-concept in this work may be far more sparse with other single-cell approaches that probe a more targeted fraction of the genome. Intriguingly, total scRNA-seq readily infers higher-order chromatin domains, owing in part to its more extensive data, but also perhaps due to the role of non-coding RNAs in targeting chromatin modifier complexes.

It has long been appreciated that nuclear RNAs play key roles in chromatin architecture. Recently, Quinodoz et al. demonstrated this with a proximity-ligation independent technique (RD-SPRITE) that readily captures and characterizes RNA and DNA interactions within the context of chromatin domains ^27,81–85^. An intriguing finding is that noncoding RNA species (ncRNA) tend to spatially localize to their own sites of transcription, likely to modulate transcription through physical means ^27^. One example of lncRNA-mediated spatial control of DNA and multi-locus gene expression regulation is the RNA–DNA interaction of *Xist* in X-chromosome inactivation of embryonic stem cells ^27^. These kinetics and structures may be somewhat transient across cell populations, and are difficult to observe in rare cells using proximity ligation assays.

With specialized assays to probe chromatin domains and interactions in single cells at kilobase resolution, it has become apparent that below the whole-chromosome scale, intra- and inter-chromosomal domain definitions are more of a statistically emergent property, averaging across many cells ^29,30,47,61^. These intra-chromosomal dynamics are also seen in domains inferred by *compartmap* from non-specialized single-cell assays. One current limitation of *compartmap* is the inability to readily distinguish observed canonical looping structures (i.e., off-diagonal increases in contact matrix densities) and inter-chromosomal interactions in a statistically principled manner. This is a focus for future work, which may be particularly important to interpret and prioritize structural variants from gene expression and chromatin accessibility data. Recent work has shown that rare SVs are abundant in human populations^86^, and can be missed even by long-read assembly-based approaches^87^. With improved prioritization schemes, the clinical benefit of long-read sequencing can be maximized relative to its cost^88^. We believe *compartmap* can serve a key role in facilitating this process for novel SVs, maximizing the value of both existing and new data, thereby accelerating discoveries.

Gene expression, methylation, and DNA accessibility are naturally linked by their underlying chromatin conformation. Recent work by Ma et al. demonstrated that what they termed the “chromatin potential” of domains of regulatory chromatin (DORCs) from simultaneously assaying chromatin accessibility and gene expression in the same cells with SHARE-seq ^59^. Similarly, SnapATAC showed that single-cell ATAC-seq (scATAC-seq) accessibility z-scores mimicked what would be observed in bulk Hi-C in GM12878 cells ^89^. Work with bulk and scRNA-seq data has shown that focal chromatin accessibility (DNase-seq signals) can be inferred from non-specialized assays at near-single-cell resolution^58^. These data demonstrate the underlying connection between gene expression and DNA accessibility through chromatin architecture.

We propose that *compartmap* provides a complementary approach to scRNAseq, scATACseq, and multi-omic single-cell assays such as SHARE-seq. Extracting the position-dependent signals from these assays reveals higher-order chromatin domains in individual cells. This offers an intuitive framework to view multiple single-cell assays through the unifying physical lens of chromatin architecture. Optimal data integration for single-cell multi-omics is an open research problem, subject to many ongoing efforts. We propose that *compartmap’s* intuitive framework of physical chromatin architecture provides immediate contextual benefits for interpreting transcriptional and epigenomic results in individual cells, whether assayed simultaneously or otherwise. At the same time, *compartmap* greatly expands the scope and scale of experiments that shed light on how chromatin architecture influences the state and fate of individual cells.

## Materials and Methods

### External datasets used

Bulk, high-resolution K562 Hi-C (bulk Hi-C) data were downloaded from GSE63525, generated by Rao et al. ^16^. Single-cell combinatorially indexed K562 Hi-C (sci-Hi-C) data were downloaded from GSE84920, generated by Ramani et al.^32^. The hg19, read-depth-normalized signal ENCODE DNase-seq bigWig track (ENCFF113ZZB) was downloaded from https://www.encodeproject.org/experiments/ENCSR000EKQ/. Lamin A/B/C TSA-seq bigWig tracks were downloaded from GSE81553. Single-cell K562 ATAC-seq data from Schep et al. were downloaded from GSE99172 ^90^. HEK293T SMART-seq total and SMART-seq3 scRNA-seq data were downloaded from GSE151334 and E-MTAB-8735, respectively. HEK293T siRNA control Hi-C fastq files (replicates 1 and 2) were downloaded from GSE44267. scRNAseq from primary AML with inv(3)(q21;q26) was downloaded from GSE116256 ^77^.

### Generation of ribo-reduced, total scRNA-seq data from K562 cells

The K562 cell line was purchased from the American Type Culture Collection (ATCC; CCL-243). Cells were cultured in RPMI 1640 medium supplemented with 10% FBS and 1% penicillin/streptomycin (Gibco). Cells were maintained at 37 °C in 5% CO_2_ and subcultured at 1:5 ratio every 3-4 d. Sorting and single-cell RNA library preparation were followed according to the Single-cell TOtal RNA Miniaturized sequencing protocol (STORM-seq; manuscript in preparation). Briefly, for live cell sorting, K562 cells were spun down at 300 × *g* for 5 min, washed in PBS (Gibco), and resuspended in 100 μL of PBS containing Zombie aqua fixable viability dye (Biolegend) at 1:1000 ratio, per 1 million cells. Cells were stained at room temperature for 15 min, quenched with 3 volumes of PBS containing 2% FBS for 3 min, washed, and resuspended in 300 μL of PBS containing 2% FBS per 1 million of cells. One, single, live cell was index-sorted into one 96-well quadrant of a 384-well plate (Eppendorf lo-bind twin-tec) containing a mixture of DPBS and shearing master mix at a final volume of 2.14 μL using a MoFlo Astrios cell sorter running Summit v6.3 (Beckman Coulter). The Astrios was equipped with a 100 μm nozzle at 25 psi. Abort mode was set to single and drop envelope to 0.5. Libraries were prepared using the STORM-seq protocol.

### Preprocessing of single-cell RNA sequencing data

K562 single-cell RNA sequencing reads (STORM-seq) were aligned using STAR (v2.7.3) to hg19 in order to be comparable to prior bulk Hi-C experiments. Aligned BAMs were then converted to bigWig format using the bamCoverage function with default parameters in deeptools (v3.4.3). BigWigs were then imported using the rtracklayer (v1.50.0) R (v4.0.3) package and mean summarized into 1-kb bins. Next, per-cell summarized bins were converted to a SingleCellExperiment (v1.12.0) object with genomic ranges added to the rowRanges slot. HEK293T SMART-seq total data were aligned with STAR (v2.7.3) to GRCh38.91 for compatibility with SMART-seq3 data. SMART-seq total HEK293T BAMs were converted to bigWigs and imported in the same manner as the K562 data. SMART-seq3 UMI counts (Smartseq3.HEK.fwdprimer.readcounts.txt) were read directly into R (v4.0.3) and converted to a SingleCellExperiment (v1.12.0) object with genomic ranges added to the rowRanges slot.

### Preprocessing of scATAC sequencing data

K562 scATAC-seq data were aligned to hg19 using bwa (v0.7.17). Next, BAMs were imported using 150-bp windows with the csaw (v1.24.3) R (v4.0.3) package. Alignments counted by csaw had to meet the following criteria: 1) non-PCR duplicate, 2) MAPQ > 20, 3) not present in the hg19 ENCODE blacklist ^91^, and 4) found on chromosomes 1-22. The background signal was determined using csaw and the aforementioned criteria but using 1-kb windows with a minimum of five reads found across the window. Foreground and background thresholding was done by fitting a mixture model to the global abundances. The end result is a background signal filtered RangedSummarizedExperiment object that can be passed to *compartmap*.

### Hi-C and sci-Hi-C chromatin domain calling

K562 .hic files were downloaded from the studies as noted above. HEK293T data were reprocessed against GRCh38.91 using Juicer (v1.5.7) and bwa (v0.7.17) with default parameters. A/B compartments from Rao et al. were computed using reads with a MAPQ > 30 and Juicer Tools (v1.14.08) at 100-kb and 1-mb resolution and normalized using the Knight-Ruiz method ^92^. HEK293T domains were called in the same manner as K562 using Juicer (v1.5.7) and Juicer Tools (v1.14.08). Sci-Hi-C data were reprocessed to hg19 single-cell 100-kb resolution contact matrices using the previously published pipeline ^32^. Chromatin domains were called by following a similar workflow previously described for single-cell HiC data ^93^. Briefly, single-cell contact matrices were converted to cooler format (cooler v0.8.10) and normalized to the lowest read depth contact matrix using the hicNormalize function in HiCExplorer (v3.5.3). Coverage normalized contact matrices were then corrected using the Knight-Ruiz method ^92^ using the hicCorrectMatrix function. Finally, chromatin domains and correlation matrices were derived using the hicPCA function. Correlation matrices were converted from .cool format to bedpe format using the R package HiCcompare (v1.12.0).

### Empirical Bayes shrinkage for bin-level compartment estimates

We reformulate the James-Stein estimator (JSE) (Equation 1) as an empirical Bayes estimator (Equation 3) as proposed in Efron et al.^94^. The multivariate normal assumption needs not be strongly enforced once the number of means exceeds 9, and it is negligible altogether once the number of means exceeds 15 ^94^.

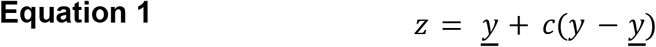

The shrunken estimate *z* for a given cell and bin is the grand mean (*<u>y</u>*) plus a shrinking factor *c* of the difference between a per-cell bin-level mean estimate (*y*) and the grand mean *<u>y</u>*. That shrinking factor, *c*, is defined below in **Equation 2**. With *k* unknown means, σ^2^ is the square of the standard deviation of bin-level estimates at a given locus across cells, and ∑ (*y* - *<u>y</u>*)^2^ is the sum of the squared deviations of per-cell bin-level estimates relative to the grand mean *<u>y</u>*.

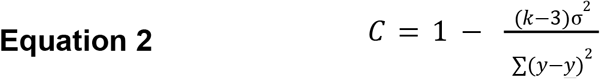

**Equation 1** can now be reformulated as the empirical Bayes estimator in **Equation 3**. The shrunken estimate *Z* now follows from the mean *m* of the prior distribution. When *k* is large, the shrinking factor *C* can assume the standard deviation of the prior on per-cell bin-level estimates (at a given locus and resolution), taking the form of an empirical Bayes estimator. However, *C* can also take the form of **Equation 2** to include the James-Stein empirical Bayes estimator for the per-cell bin-level estimates, provided *k* is greater than or equal to 4. We note that *m* need not be a single, global scalar, but can with mild constraints be a vector of group-wise means, without loss of generality.

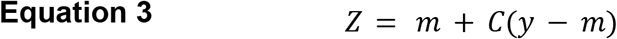

Various iterative imputation and semi-supervised clustering approaches follow naturally from **Equation 3** when per-group mean and membership estimates are obtained via expectation-maximization. Under the mild constraint that any given group is of sufficient size (*N* >= 15), the previous remarks from Efron et al.94 hold at the group level, and we can thus expect monotonic convergence to a local minimum error, even in the presence of data-adaptive shrinkage (see ^95^ for further discussion).

### Inference of higher-order chromatin in single cells with *compartmap*

Preprocessed single-cell assay objects that contain genomic ranges can be used as input to *compartmap*. First, features are TF-IDF transformed. Next, single-cell or group-level higher-order chromatin domains are computed using the scCompartments function in *compartmap*. Within this function, multiple steps are performed. Bootstrap global means are first taken to speed up computation time. Briefly, *N* bootstrap genome-wide global means are drawn from the data. Following the initial computation of chromatin domains in a single-cell (described below), the procedure is repeated for each bootstrap global mean. The results are then tallied into “open” or “closed” states using the signs of the first right singular vector, followed by computing Aggresti-Coull confidence intervals for each genomic bin ^64^. Given a confidence estimate for each genomic bin, enabling users to invert the bin (open -> closed or closed -> open) if it exceeds some threshold, or to use it as a way to weight the initial estimates. To compute single-cell or group-level higher-order chromatin domains, the TF-IDF transformed estimates are shrunken towards either a global or targeted mean using the JSE described above. Next, Pearson correlations are computed to either the global/targeted mean in the case of single-cell domains or to within-group pairwise comparisons in the case of group-level. Then, SVD is performed on the correlation matrix using IRLBA as implemented in the BiocSingular package (v1.6.0) and retaining the first right singular vector. Due to the arbitrary assignment of signs to the eigenvector, we attempt to ensure consistency of signs by checking that the right singular vector has a positive correlation to the column sums of the binned, shrunken correlation matrix, changing signs if necessary as previously described ^56^. Finally, the first right singular vector is smoothed using a windowed smoother, as previously described^56^.

### Hi-C and sci-Hi-C correlation matrix generation

K562 and HEK293T bulk Hi-C Pearson correlation matrices were generated using Juicer Tools (v1.14.08) at 1-mb resolution from Juicer-derived (v1.5.7) .hic files with a MAPQ > 30 and KR normalized. K562 sci-Hi-C Pearson correlation matrices at 1-mb resolution were generated as described above with HiCExplorer (v3.5.3) and HiCcompare (v1.12.0).

### Generation of scRNA-seq and scATAC-seq correlation matrices

Raw, group-level correlation matrices for K562 and HEK293T scRNA-seq or scATAC-seq were generated as described above. However, the raw correlation matrices are noisy and were denoised using random matrix theory. Briefly, binned, shrunken group-level single-cell correlation matrices were computed at 1 Mb resolution following preprocessing described above for the respective assays. We take advantage of the correlation structure by fitting a Marchenko-Pastur density to the eigenvalues of the input correlation matrix, which in effect, these eigenvalues represent the higher-order chromatin domains. The denoising occurs by setting a lower-bound on the fitted eigenvalues and retaining those above the computed threshold. The function utilized is wrapped in *compartmap* and implemented in the R package covmat (v1.1).

## Acknowledgements

We thank all members of the Triche and Shen labs, in addition to Peter Jones, Elana Fertig, Genevieve Stein-O’Brien, Peter Laird, Patrick Grohar, Matt Steensma, Carrie Graveel, Maggie Chasse, Josh Jang, Brittany Carpenter, Zach Madaj, Emily Wolfrum, Jamie Grit, Brad Dickson, and Andrew Pospisilik for insightful suggestions and thoughtful discussions. We would like to acknowledge the Van Andel scientific computing team for providing computational resources for this work, and David Nadziejka for technical editing support. This work was supported by the Van Andel Institute, the Michelle Lunn Hope Foundation, the Folz Family Foundation, an anonymous VAI donor’s bequest to the laboratory of T.T., and an NIH R37 awarded to H.S. (R37CA230748).

## Author contributions

B.K.J., H.S., and T.T. conceived the project. B.K.J and T.T. implemented *compartmap*, with substantial previously unpublished contributions from K.D.H. and J.P.F. in early iterations. B.K.J. led the data analysis. B.K.J., H.S., and T.T. wrote the manuscript. All authors read and approved the manuscript.

## Data availability

K562 STORM-seq data are available from GSEXXXXX.

Primary inv(3) AML scRNAseq data is available from GSE116256 (subject AML328). Intermediate data can be downloaded from Zenodo, DOI: 10.5281/zenodo.4763579.

## Software availability

*Compartmap* is available as a Bioconductor package: https://www.bioconductor.org/packages/devel/bioc/html/compartmap.html

