## Supplemental figures for "*Compartmap* enables inference of higher-order chromatin structure in individual cells from scRNA-seq and scATAC-seq"

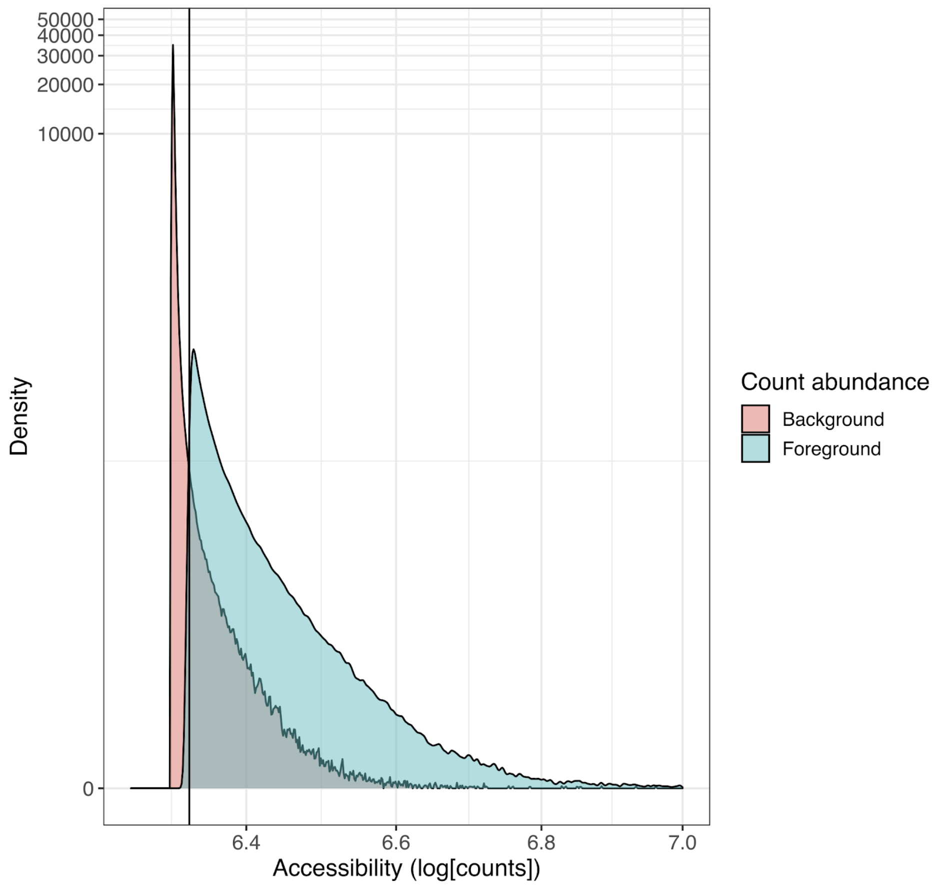


**Figure S1. Mixture model filtering of scATAC-seq windows.** As an initial quality control procedure, a two-component mixture model was fit to the windowed scATAC-seq $\log_2$ count data to filter out low-abundance background (pink color) across single cells while preserving accessibility signals (foreground - blue). Background was determined as windows less than the $\log_2$ count cutoff (black vertical line) separating the two populations (pink - background; blue - foreground).

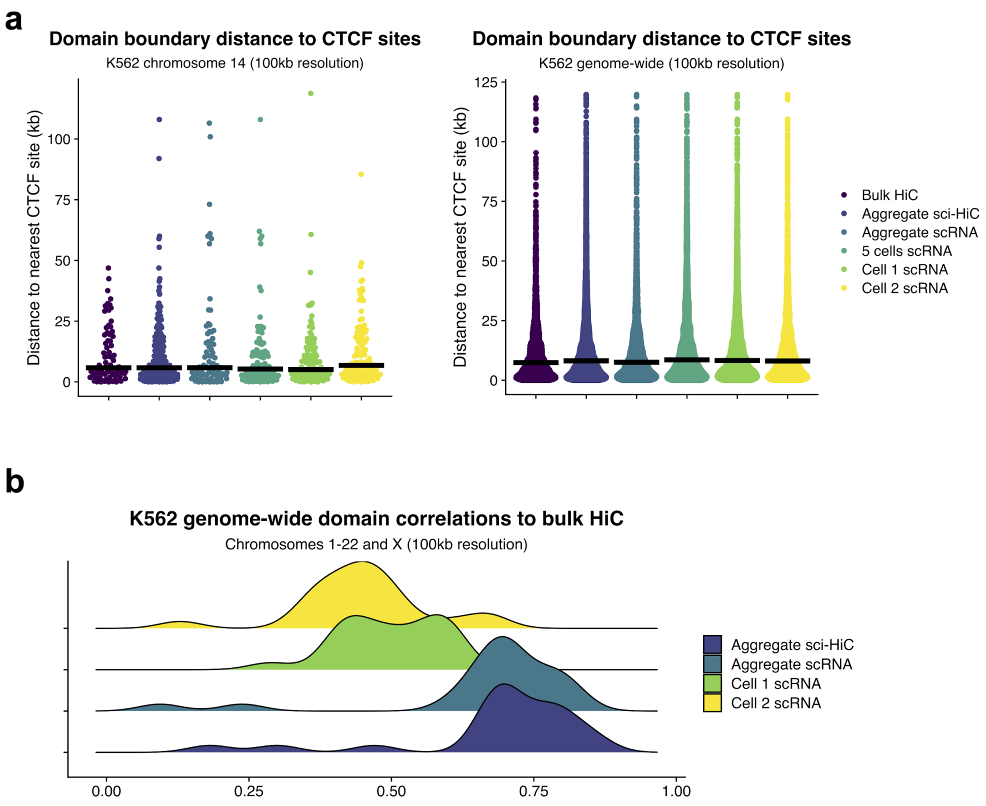


**Figure S2. *Compartmap*-inferred higher-order chromatin domains from K562 total scRNA-seq preserve domain boundary distances to CTCF sites and correlates to bulk Hi-C similar to sci-Hi-C.** (a) *Compartmap*-inferred chromatin domain boundary distances to nearest CTCF sites from K562 scRNA-seq are consistent relative to boundary distances derived from chromatin-capture-based methods like Hi-C and sci-Hi-C. The hg19 ENCODE v3 CTCF ChIP-seq sites were downloaded from the UCSC Genome Browser for determining distance to the nearest CTCF site. (b) Correlations of chromatin domain eigenvalues from sci-Hi-C and *compartmap*-inferred chromatin domains from scRNA-seq to bulk Hi-C at 100-kb resolution genome-wide. Aggregate sci-Hi-C, scRNA-seq, and scRNA-seq cells 1 and 2 represent the same cell populations as shown in **Figure 2a**.

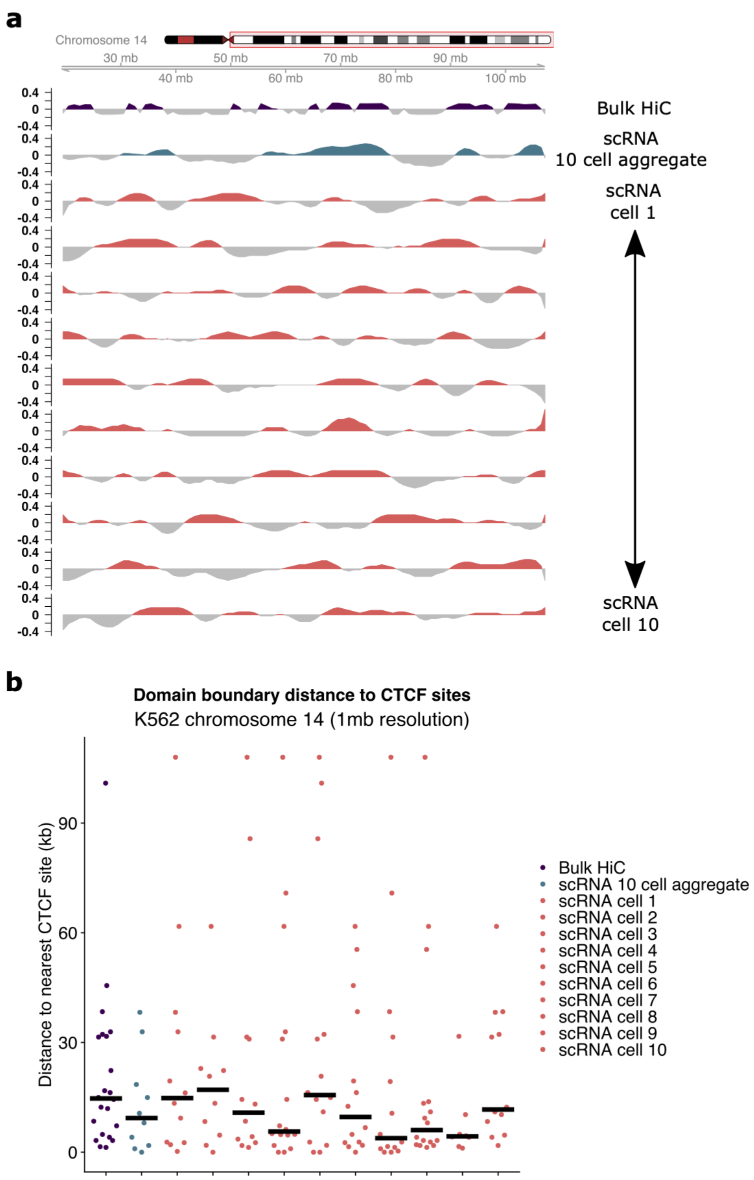


**Figure S3. Single-cell K562 chromatin domains are heterogenous but in aggregate provide a similar view as bulk Hi-C while maintaining coherent distances to CTCF sites.** **(a)** *Compartmap* reconstruction of higher-order chromatin domains (1-mb resolution) in 10 arbitrarily chosen single cells (original population *N* = 70 as shown in **Figure 2a**) demonstrate heterogeneous chromatin domains relative to bulk Hi-C. However, in aggregate, the 10 cells from scRNA-seq largely reconstruct features observed in bulk Hi-C. Positive eigenvalues (non-gray colors) represent “open” chromatin domains. Negative eigenvalues (gray color portion of the track) represents “closed” chromatin domains. **(b)** Despite the heterogeneity observed in single-cell inferred higher-order chromatin, the domain boundaries maintain similar distances to CTCF sites. ENCODE V3 CTCF sites were downloaded from the UCSC genome browser for hg19.

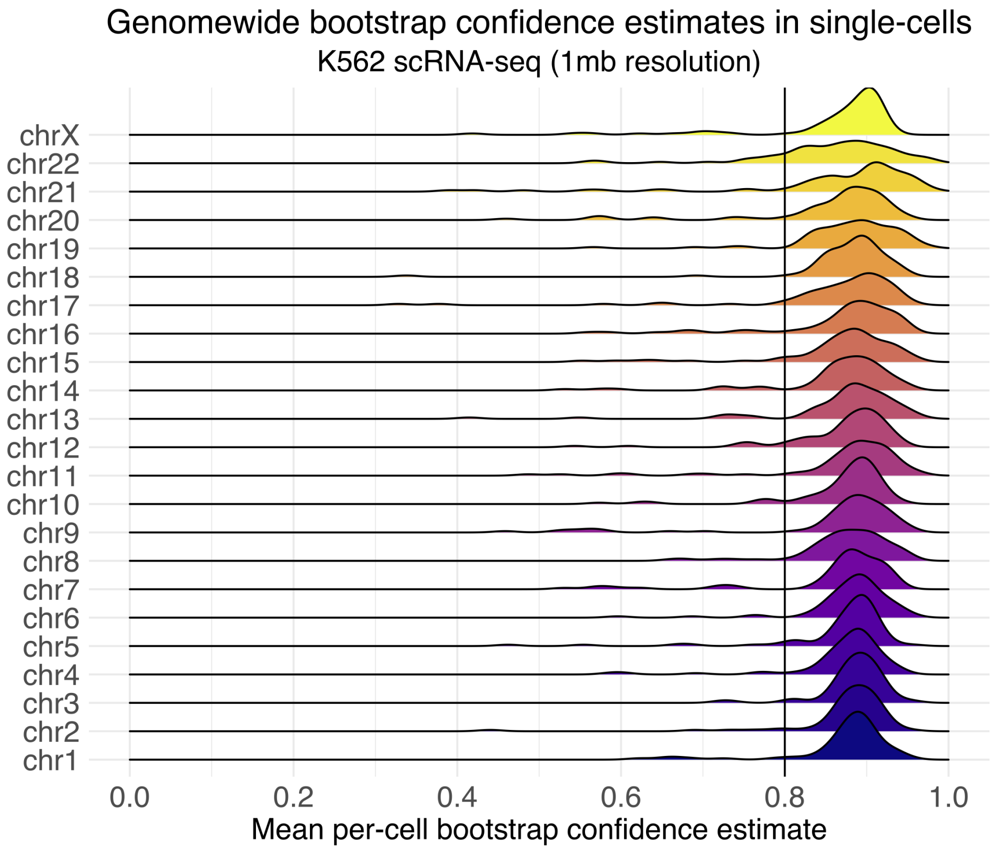


**Figure S4. Genome-wide mean per-cell bootstrap confidence estimates for scRNA-seq K562 inferred chromatin domains demonstrate high-confidence compartment-scale inference in single cells with *compartmap*.** The density of single-cell, per-chromosome mean bootstrap confidence estimates (100 bootstraps) of inferred chromatin domains (1-mb resolution) from K562 scRNA-seq demonstrates that *compartmap* is able to confidently reconstruct (>80%; black line) higher-order chromatin in single cells. Additionally, this metric provides a mechanism to directly assess variability and confidence across user-defined genomic scales when comparing group-level or single-cell chromatin domains.

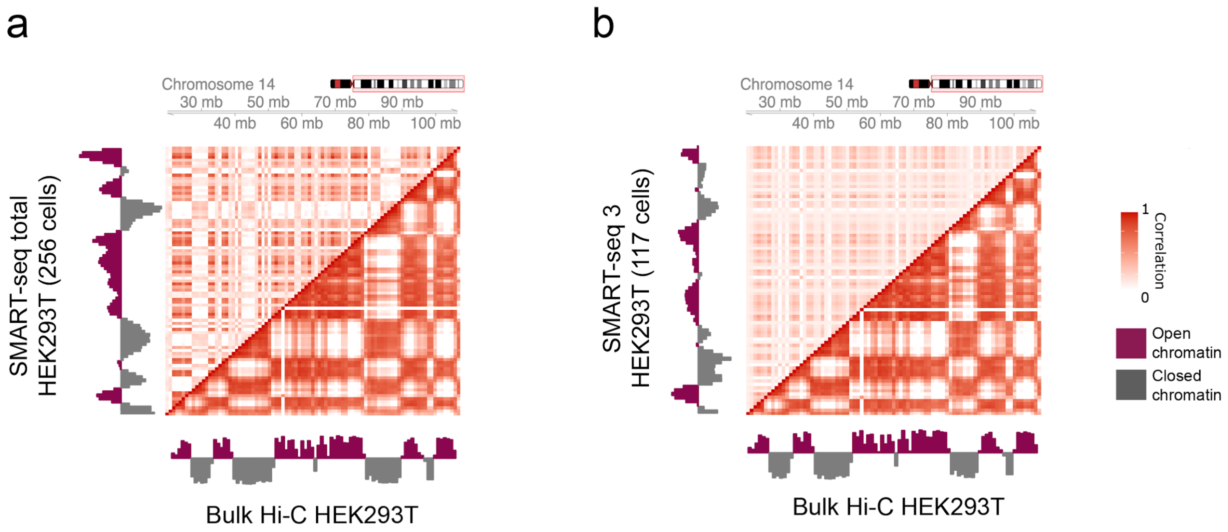


**Figure S5. *Compartmap* infers higher-order chromatin across scRNA-seq platforms and cell types.** Comparison of group level correlation matrices of chromosome 14 at 1-mb resolution in HEK293T cells show similar resolution and structure between (a) SMART-seq total scRNA-seq ($N$ = 256 cells) and bulk Hi-C and (b) SMART-seq 3 ($N$ = 117 cells) and bulk Hi-C. scRNA-seq correlation matrices were generated using *compartmap*. Bulk Hi-C correlation matrices were computed from normalized contact matrices. Coloration represents low (white) to high (red) within-platform correlations and corresponding “open” (maroon) or “closed” (gray) higher-order chromatin states.
